# E41K Mutation Activates Bruton’s Tyrosine Kinase by Stabilizing an Inositol Hexakisphosphate dependent Invisible Dimer

**DOI:** 10.1101/2024.01.22.576617

**Authors:** Subhankar Chowdhury, Manas Pratim Chakraborty, Swarnendu Roy, Kaustav Gangopadhyay, Rahul Das

## Abstract

Bruton’s tyrosine kinase (BTK) regulates diverse cellular signaling of the innate and adaptive immune system in response to microbial pathogens. Downregulation or constitutive activation of BTK is reported in patients with autoimmune diseases or various B-cell leukemias. BTK is a multidomain protein tyrosine kinase that adopts an Src-like autoinhibited conformation maintained by the interaction between the kinase and PH-TH domains. The PH-TH domain plays a central role in regulating BTK function. The BTK is activated by binding to PIP_3_ at the plasma membrane upon stimulation by the B-cell receptor (BCR). The PIP_3_ binding allows dimerization of the PH-TH domain and subsequent transphosphorylation of the activation loop. Alternatively, a recent study shows that the multivalent T-cell-independent (TI) antigen induces BCR response by activating BTK independently of PIP_3_ binding. It was proposed that a transiently stable IP_6_-dependent PH-TH dimer may activate BTK during BCR activation by the TI antigens. However, no IP6-dependent PH-TH dimer has been identified yet. Here, we investigated a constitutively active PH-TH mutant (E41K) to determine if the elusive IP_6_-dependent PH-TH dimer exists. We showed that the constitutively active E41K mutation activates BTK by stabilizing the IP_6_-dependent PH-TH dimer. We observed that a downregulating mutation in the PH-TH domain (R28H) linked to X-linked agammaglobulinemia impairs BTK activation at the membrane and in the cytosol by preventing PH-TH dimerization. We conclude that the IP_6_ dynamically remodels the BTK active fraction between the membrane and cytoplasm. Stimulating with IP_6_ increases the cytosolic fraction of the activated BTK.

## Introduction

Bruton’s tyrosine kinase (BTK) is an indispensable signaling module of innate and adaptive immune response against pathogens. BTK downregulation in X-linked agammaglobulinemia (XLA) patients causes recurring bacterial and viral infection (1–4). On the other hand, constitutive activation of BTK is often linked to several forms of B-cell leukemias and lymphomas like chronic lymphocytic leukemia (B-CLL) (5), and diffuse large B cell lymphoma (ABC-DLBCL) (6), suggesting a critical regulatory function of BTK in B-cell development and proliferation (7). Thus, BTK has been an important pharmacological target for the treatment of various B-cell malignancies (8–10).

In macrophages, a component of innate immunity, BTK phosphorylates the Toll-like receptors (TLR) to initiate an antiviral response (11). In B-cells, a component of adaptive immunity, BTK is essential for connecting the B-cell antigen receptor (BCR) activation to the downstream calcium flux (12–14). BTK is a Tec family kinase (15) that shares a homologous structural architecture with the SRC kinases, where the kinase domain is flanked by two N-terminal SRC homology domains, SH3 and SH2 (Figure 1A) (16–18). BTK has an additional lipid binding pleckstrin homology (PH) domain fused to the Tec homology (TH) domain at the N-terminal of the SH3 domain (19,20). Unlike Src kinase, where the autoinhibited conformation is stabilized by the C-terminal phosphotyrosine residue (16–18), in BTK, the PH-TH domain functions as the negative regulator (21–23). In the autoinhibited state, the SH3 and SH2 domain assemble at the back of the kinase domain (Figure 1B) (16–18,22). The autoinhibited structure is maintained by the interaction of the PH-TH domain and the kinase domain, blocking the activation loop face of the kinase domain, which occludes the lipid binding to the PH-TH domain (22,24,25).

**Figure 1.**
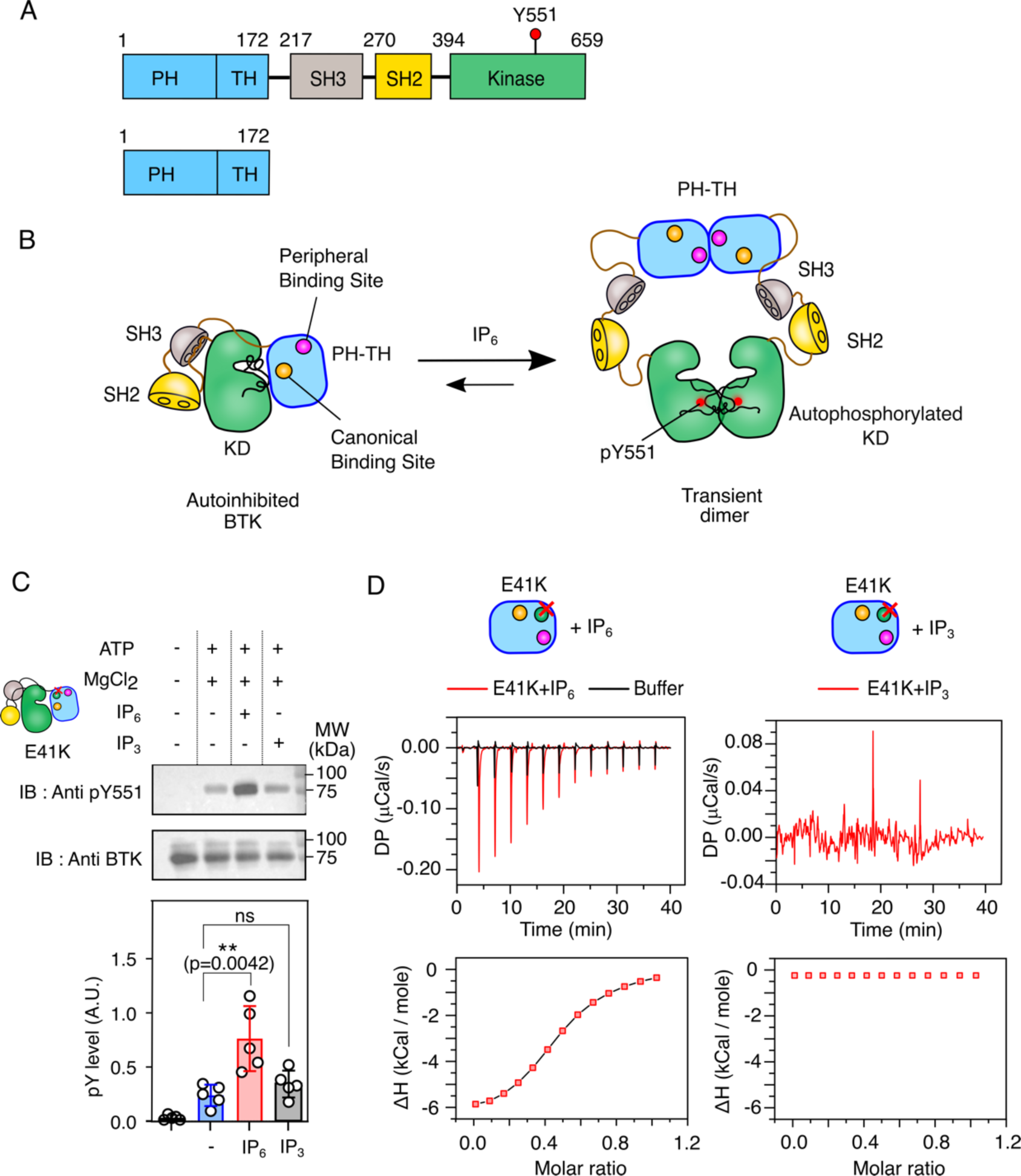
Activation of BTK E41K mutant by Inositol Hexakisphosphate (IP_6_) (A) Schematic representation of domain architecture of Bruton’s Tyrosine Kinase and PH-TH domain used in this study. (B) Mechanism of Inositol Hexakisphosphate (IP_6_) dependent activation of Bruton’s Tyrosine Kinase (22). (C) Top panel: representative immunoblot of Y551 autophosphorylation in the full-length BTK E41K (2μM) mutant with 100 μM IP_6_ or IP_3._ Bottom panel: densitometric analysis of the immunoblots of BTK E41K mutant activated with IP_6_ or IP_3_. The error bar represents standard deviations from five independent experiments. Data are presented as mean values ± SD from five independent experiments. Data analyses were performed using GraphPad Prism version 9.5.1. An unpaired two-tailed t-test was used to calculate significance. (D) Top panel: Isothermal Titration Calorimetric (ITC) measurement of IP_6_ (left) or IP_3_ (right) binding to E41K mutant of an isolated PH-TH domain. For each titration, 25μM of PH-TH^E41K^ was titrated with 137 μM of ligand. Bottom panel: solid line represents the two-site independent binding fitting of the ITC titration of IP_6_ and indicated PH-TH domain. The experimental points are indicated in red square. The ITC titration of IP_3_ and PH-TH^E41K^ domain could not be fit to any model. The plots were generated using Origin Pro 2020b.

Antigen binding to BCR activates BTK by recruiting kinase to the plasma membrane (Figure 1SA) (26,27). The localization of the BTK to the membrane is mediated by the binding between the phosphatidylinositol 3–5-triphosphate (PIP_3_) and canonical lipid-binding site of the PH-TH domain (Figure 1SA) (4,22,28,29). Binding to PIP_3_ removes the inhibitory interaction allowing BTK to dimerize (28,30) and rapidly phosphorylate the tyrosine residue in the activation loop (13,31,32). The PIP_3_ induces dimerization of the PH-TH domain (called Saraste dimer) where hydrophobic dimer interfaced (called Saraste interface) is constituted by α1-helix, β1 and β3-β4 loop (22,28,29). At the membrane, the PH-TH dimer is stabilized by a second PIP_3_ binding to the peripheral lipid binding site (Figure 1SA), which imparts a switch-like property to the kinase (30).

Alternatively, recent studies showed that inositol hexakisphosphate (IP_6_) is required for T-cell independent BCR response against bacterial and viral infection (33). Deficiency of inositol polyphosphate multikinase in mice, the enzyme that generates the precursor for IP_6_ biosynthesis from inositol tetrakisphosphate (IP_4_) (34), impairs BTK activity and attenuates calcium signaling in B-cells. The IP_6_ was previously shown to induce BTK activation in solution by the same PH-TH Saraste dimer, where two molecules of IP_6_ bind to the canonical site and peripheral sites, respectively (Figure 1B) (22), suggesting IP_6_ activates BTK independently of its membrane localization. Mutations that impair IP_6_ binding to the peripheral sites inhibit BTK activation in the solution. However, no PH-TH dimer in solution has yet been determined. It was speculated that the PH-TH domain might form a transiently stable dimer in the presence of IP_6_ (22). Therefore, the lack of evidence for an IP_6_-dependent BTK dimer makes the functional relevance of T-cell indigent BCR response to pathogens uncertain.

In the present study, we focused on an E41K mutant of the PH-TH domain that activates BTK constitutively (35) and binds to IP_6_ with a higher affinity compared to the wild-type (4). We observed that unlike wild-type BTK (22), the E41K mutant is activated by IP_6_ in solution by stabilizing a PH-TH dimer, which was otherwise invisible by most biophysical techniques due to the transient nature of the dimer. Our mutation studies suggest that the PH-TH dimerization is mediated by an alternate dimer interface, where an IP_6_ molecule holds two lateral PH-TH domains together. The R28H mutation found in the canonical lipid-binding site of the PH-TH domain in XLA patients (4,36) prevents IP_6_-dependent PH-TH dimerization and inhibits BTK activation. In cells, higher IP_6_ concentration increases the cytosolic fraction of activated BTK. Finally, we presented a mechanism explaining how IP_6_ remodels the equilibrium between the membrane fraction and cytosolic fraction of BTK in the cell.

## Results and Discussion

### IP_6_ activates BTK E41K mutant in solution

The E41K mutation is located at the β3-β4 loop of the PH-TH domain (Figure 2A). The lysine residue does not make direct contact with the IP_4_ molecule at the canonical binding site (37). Instead, the lysine creates an overall positively charged environment, allowing the second IP_4_ molecule to bind, leading to spontaneous membrane recruitment of the BTK, autophosphorylates Y551 at the activation loop, and enhanced Ca^2+^ signaling (35,37,38). The high structural homology (RMSD of 0.43 Å) between the wild-type BTK (PDB ID: 1bwn) (37) and the E41K mutant (PDB ID: 4y94) (22) does not explain the observed functional difference (Figure 1SB). However, a higher binding affinity of the E41K mutant for IP_6_ (4) suggests that the mutation may stabilize a PH-TH dimer. To find out if IP_6_ also activates BTK carrying an E41K mutation in solution, we purified the full-length BTK (denoted as BTK^E41K^) and an isolated PH-TH domain with E41K mutant (denoted as PH-TH^E41K^) and determined the phosphorylation of Y551 and binding of IP_6_, respectively (Figure 1C). We observed that in the absence of any ligand, the BTK^E41K^ spontaneously phosphorylated at Y551. Whereas the addition of IP_6_ significantly increases the Y551 phosphorylation compared to the IP_3_ and the unligated state. As reported previously, IP_3_ does not induce phosphorylation of Y551 because the PH-TH^E41K^ domain does not bind to IP_3_ (Figure 1D and Table S1) (4). On the other hand, PH-TH^E41K^ binds to IP_6_ with two distinct dissociation constants (*K_d1_*= 152 ± 18 nM and *K_d2_*= 1.3 ± 0.4 µM) when the ITC data was fitted to two-site independent binding model (Figure 1D and Table S1). We observed that the *K_d1_* and *K_d2_* represent the strong and the weak IP_6_ binding sites, respectively, which may correspond to the canonical and peripheral lipid binding sites of the PH-TH domain, respectively (22). We asked if the improved activation of BTK is due to the greater stability of the IP_6_-dependent dimer in the solution.

**Figure 2.**
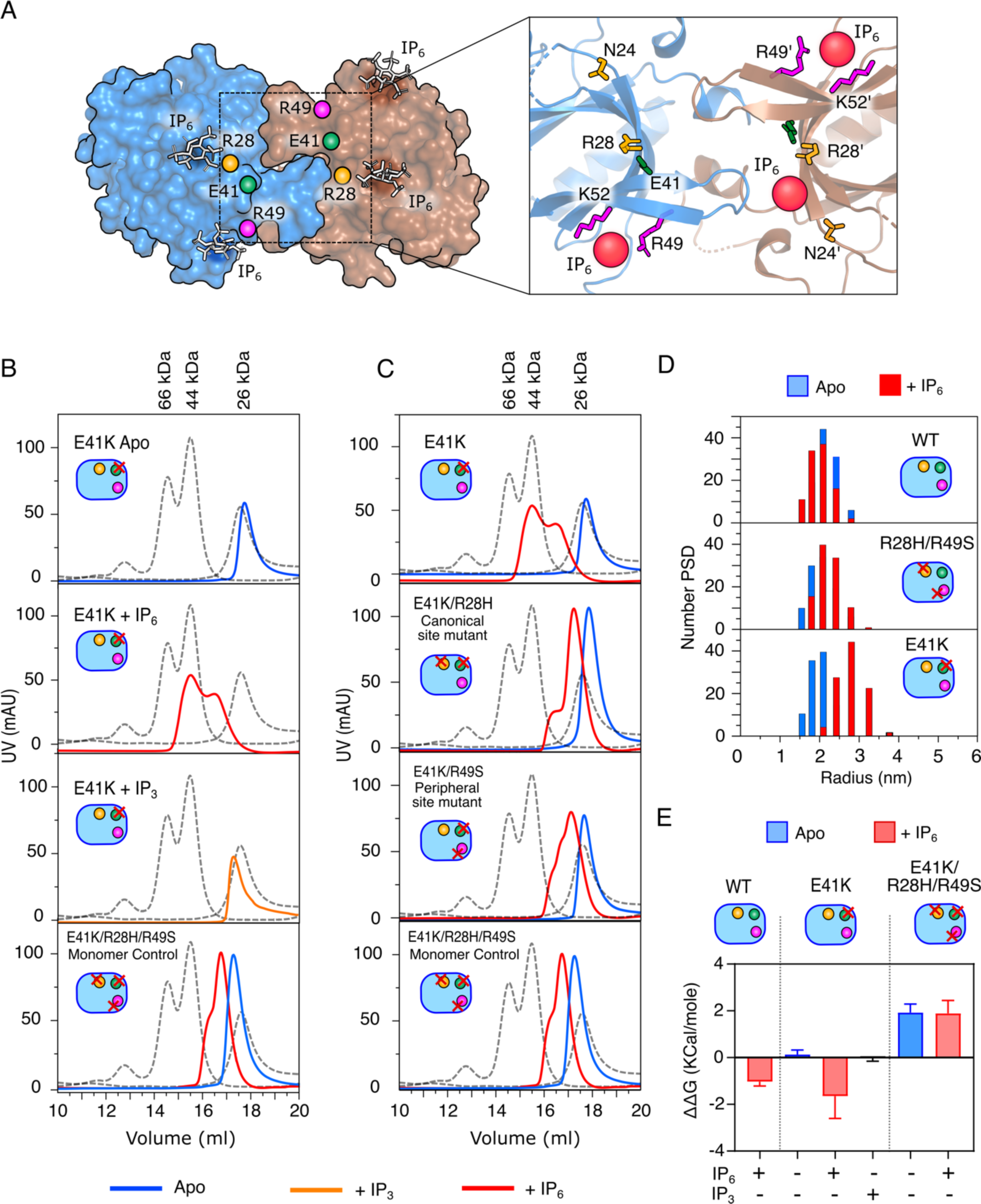
Probing the IP_6_-dependent PH-TH dimer in solution. (A) The space-filled model of BTK PH-TH Saraste dimer in complex with IP_6_ (PDB 4Y94). The canonical and peripheral ligand binding site is shown in the inset. (B and C) Representative gel-filtration elution profiles of indicated constructs of PH-TH domain in the presence (300 μM) or absence of IP_6_ or IP_3_. The triple mutant of the PH-TH domain (E41K/R28H/R49S) is used as monomer control. The dotted line represents the elution profiles of standard protein mixtures containing BSA (66 kDa), Ovalbumin (44 kDa), and ULP1 (26 kDa). The plots were generated by XMGRACE Ver 5.1.25. (D) Dynamic Light Scattering measurement of indicated constructs of PH-TH domain in the presence (red bar) or absence (blue bar) of IP_6_. The plots were generated using Origin Pro 2020b. (E) ΔΔG_unfolding_ of Apo and ligand-bound BTK PH-TH constructs derived from the thermal denaturation profile of the Circular Dichroism spectrum. Data are presented as mean values ± SD from three independent experiments. The plots were generated using GraphPad Prism version 9.5.1.

### IP_6_ promotes PH-TH dimerization

In the crystal structure of the PH-TH domain in complex with IP_6_, the Saraste-dimer interface is comprised of antiparallel interaction between the adjacent β3-β4 loop and α1-helix of each PH-TH domain (Figure 2A) (22). However, due to the transient nature of the hydrophobic interaction at the Saraste-dimer interface, no IP_6_-dependent PH-TH dimer is yet being detected in the solution. Since PH-TH^E41K^ binds IP_6_ with a stronger affinity compared to the wild-type (4), we speculate if the activation of PH-TH^E41K^ is due to the formation of a stable IP_6_-dependent dimer. We probed the PH-TH^E41K^ dimer from the mobility of the protein in a gel-filtration column and from dynamic light scattering (DLS) measurements (Figure 2). In our assays, we used double mutant (R28H and R49S at the canonical and peripheral site, respectively) of the PH-TH domain made in the wild-type or E41K background as a monomer control. The anticipated molecular weight and theoretical hydrodynamic radius of the PH-TH dimer are 38 kDa and 3.0 nm, respectively (Figure S2B). As reported previously, we could not detect IP_6_-dependent dimers for the wild-type PH-TH domain by gel-filtration assay and DLS experiment (Figure S2 A-C and Figure 2D). We observed a minor shift in the retention time for the IP_6_ bound PH-TH domain compared to the *apo*-state, and the hydrodynamic radius (2.18 ± 0.10 nm) was close to a monomer (Figure S2A-C and Figure 2D).

On the contrary, the PH-TH^E41K^ in the presence of IP_6_ migrated close to a dimer (44 kDa) species on a gel-filtration column (Figure 2B). The hydrodynamic radius of 3.14 ± 0.48 nm suggests that the PH-TH^E41K^ in the presence of IP_6_ forms a stable homodimer (Figure 2D, Figure 2B-C). We noted that PH-TH^E41K^ does not dimerize in the *apo*-state or in the presence of IP_3_ (Figure 2B). Comparison of change in Gibb’s free energy for unfolding (ΔΔG_unfolding_) suggests that the PH-TH^E41K^ in complex with IP_6_ has relatively higher stability compared to IP_3_ or wild-type PH-TH domain (Figure 2E, S2D and Table S2). Together our data suggests that the activating mutant or ligand stabilizes the PH-TH domain structure compared to inactivating mutants or ligands (Figure 2E) (29).

### Activation of BTK E41K mutant is mediated by the IP_6_-dependent PH-TH dimer

The β3-β4 segment crosstalk with the canonical lipid binding site, peripheral lipid binding site and the Saraste dimer interface. Recent studies suggest that PIP_3_ binding to the canonical and peripheral lipid binding is important for membrane docking and dimerization of BTK (29,30). Mutation, such as R28H found in XLA patients, at canonical lipid binding pocket impairs PIP_3_ binding and down-regulates BTK signaling (39). We next investigate whether the down-regulation of BTK in XLA mutant found at the PH-TH domain is due to impaired dimerization.

We begin with characterizing IP_6_ binding and dimerization of the R28H mutant (at the canonical site) and R49S mutant (at the peripheral site) created in the PH-TH^E41K^ background (Figure 2C, 3A, and Table S1). We also compared the ability to autophosphorylate full-length BTK E41K mutant (BTK^E41K^) bearing a mutation in the canonical (R28H) or peripheral (R49S) lipid binding site (Figure 3B-C). We observed that the R28H mutation impairs IP_6_ binding to the canonical site but retains the weak IP_6_ binding to the peripheral lipid binding site with *K_d_* of 2.4 ± 0.15 µM (Figure 3A and Table S1). The R49S mutation in the peripheral lipid binding site impairs IP_6_ binding at the peripheral site but retains the strong IP_6_ binding to the canonical lipid binding site with *K_d_* of 622 ± 77 nM. However, in the gel-filtration assay, we observed that none of the mutants dimerize in the presence of IP_6_ and migrate along with the monomer control (Figure 2C). As anticipated, impaired PH-TH dimerization impairs IP_6_ dependent autophosphorylation of BTK^E41K^ R28H or R49S mutant (Figure 3B-C). Our data shows that the XLA mutation (R28H) and mutation at the peripheral lipid binding site (R49S) impair BTK function by destabilizing the Saraste dimer interface, suggesting an allosteric crosstalk between the two lipid binding pockets and the dimer interface.

**Figure 3.**
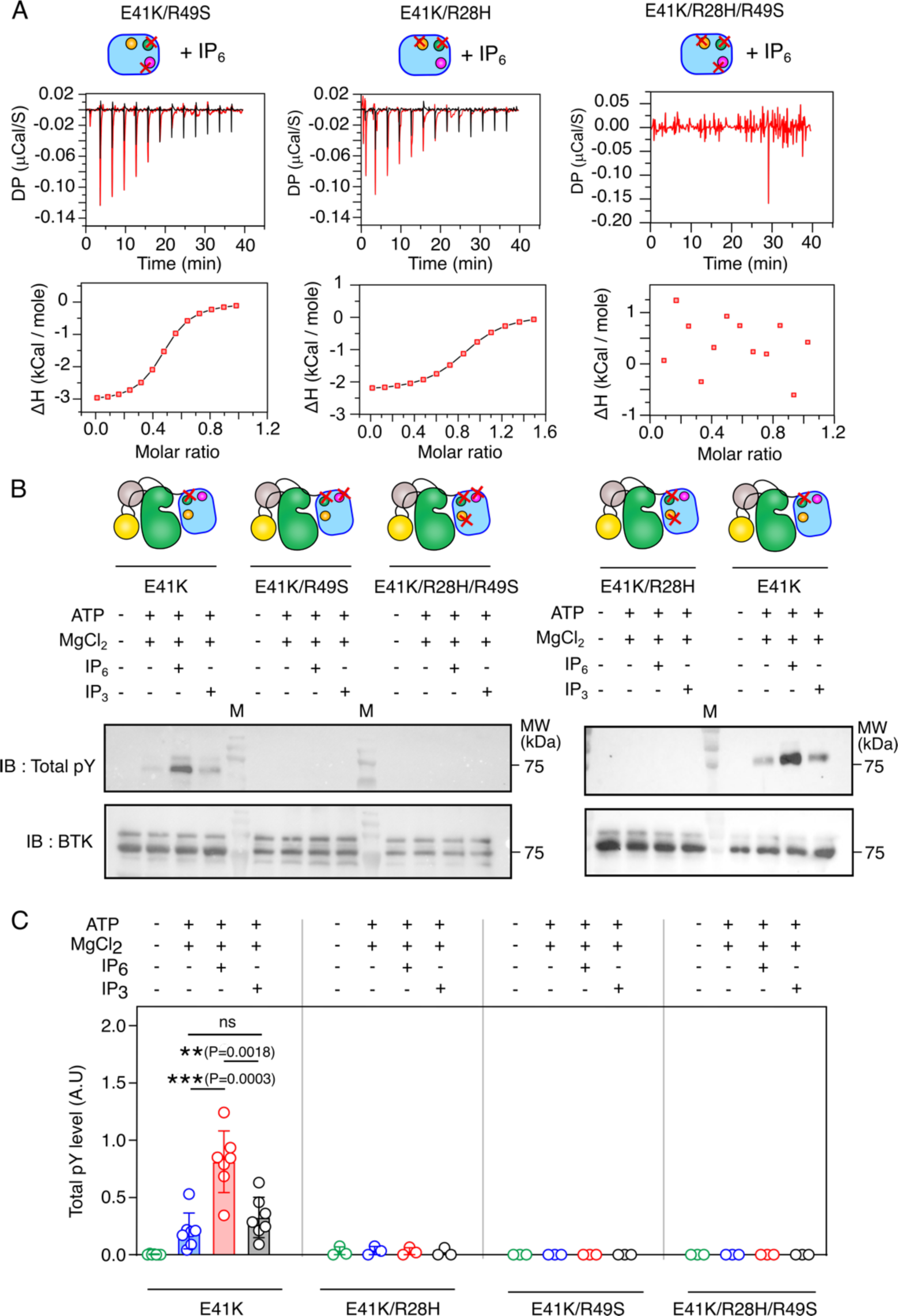
Functional analysis of IP_6_ mediated activation of BTK E41K mutant. (A) ITC titration of IP_6_ to the indicated mutants of the PH-TH domain of BTK. For each titration, 25μM of PH-TH^E41K^ was titrated with 137 μM of ligand. Top panel: red lines indicate protein and ligand titration, and the black line represents buffer-to-buffer titration. Bottom panel: solid line represents the one-site binding fitting of the ITC titration of IP_6_ and indicated PH-TH domain. The experimental points are indicated in red square. The ITC titration of IP_3_ and PH-TH^E41K^ domain could not be fit to any model. The plots were generated using Origin Pro 2020b. (B) Representative immunoblot of indicated mutants of full-length BTK with 100 μM IP_6._ The level of autophosphorylation is determined with a total anti-phosphotyrosine antibody. (C) Densitometric analysis of the immunoblots in panel B. Data are presented as mean values ± SD from 5-7 independent experiments for E41K and 3 independent experiments for indicated double and triple mutants of PH-TH domain (E41K/R28H, E41K/R49S, E41K/R28H/R49S). An unpaired two-tailed t-test was used to calculate significance. Data analyses were performed using GraphPad Prism version 9.5.1.

### E41K mutation promotes spontaneous localization of BTK to the plasma membrane

In B-cells, BTK is activated by localizing the kinase to the membrane following antigen binding to the BCR (26,40). The E41K mutation spontaneously localizes the kinase to the plasma membrane when expressed in NIH 3T3 cells and trans phosphorylate Y551 in the activation loop (13,35,38). At the membrane, the binding to PIP_3_ at the canonical and peripheral lipid binding sites activates BTK by stabilizing the Saraste dimer (30). Recently, IP_6_ was shown to regulate BTK function during BCR response to T-cell independent antigens (33). We wonder about how IP_6_ may remodel the BTK activation profile at the membrane.

To study the effect of IP_6_ on the activation profile of BTK, we transiently transfected wild-type and E41K mutant (called BTK^E41K^) of BTK-tagged mCherry to the CHO cells and determined the membrane localization (Figure 4A-B). The plasma membrane was marked with Wheat Germ Agglutinin (WGA) fused to Alexa 633 (41). The activation of BTK was determined with a specific anti-pY551 antibody and then imaged by labeling with a FITC-tagged secondary antibody (Figure 4). The membrane localization of BTK^E41K^ was estimated from the quantification of Pearson’s colocalization coefficient (PCC) of mCherry and Alexa 633 (42). A higher PCC value indicates higher membrane localization. We used a triple mutant (E41K, R28H, and R49S) of BTK as a monomer control (negative control) (Figure S2 B-D). We observed that a major fraction of wild-type BTK remains in the cytosol (Figure 4A and S3D). In contrast, a significant portion of the BTK^E41K^ localized to the plasma membrane (Figure 4B). Consistent with our *in-vitro* experiments (Figures 2 and 3), we observed that the mutation at the peripheral lipid binding site (R49S), made in the background of BTK^E41K^, localized to the membrane (Figure 4C). However, the XLA mutation (R28H) impairs the membrane localization of BTK^E41K^ (Figure 4D). As shown previously, we also observed that only the BTK^E41K^ spontaneously phosphorylated Y551 at the plasma membrane (Figure 4B and S3A), but none of the other mutants were active (13,35,38). These data suggest that the BTK in the CHO cells is mainly recruited to the membrane by binding PIP_3_ to the canonical lipid-binding site of the PH-TH domain (29,30,38). Binding of PIP_3_ to the peripheral lipid binding site is required for the transphosphorylation of BTK (22,30). We asked if BTK^E41K^ is activated only at the plasma membrane or if the cytoplasmic fraction of BTK^E41K^ could be stimulated by IP_6_.

**Figure 4.**
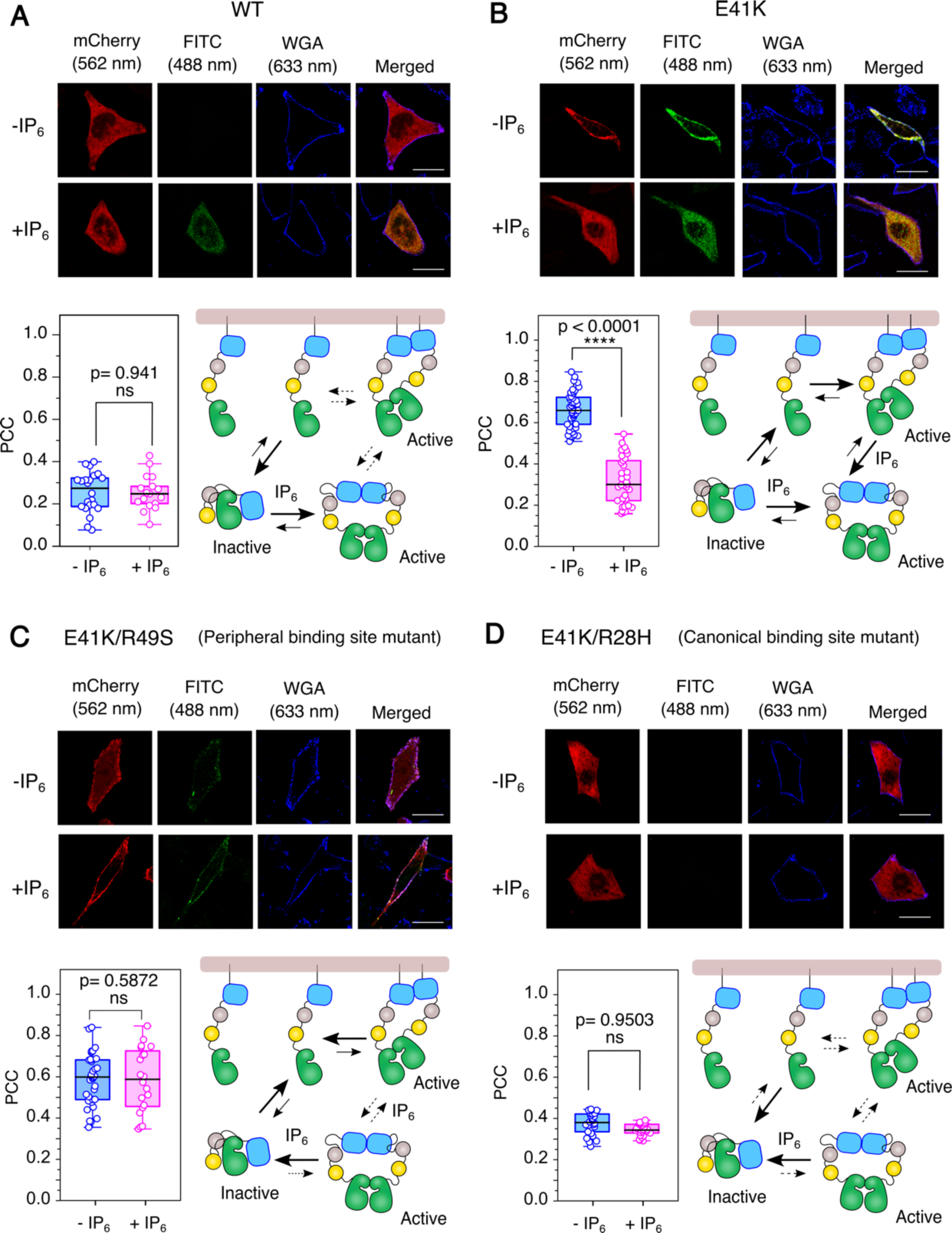
IP_6_-dependent activation of BTK in CHO cell line. (A-D) Confocal images of transiently transfected BTK constructs fused to mCherry in CHO cell lines. The BTK expression level is shown in red (λ_ex_= 552 nm, λ_em_= 586-651 nm), and the phosphorylation status is shown in green (λ_ex_= 488 nm, λ_em_= 505-531 nm). The blue represents the plasma membrane stained with Wheat Germ Agglutinin (WGA) fused to Alexa 633 (λ_ex_= 633 nm, λ_em_= 647-692 nm). In each panel, the first column is the expression level of the indicated construct of BTK-mCherry. The second column represents the phosphorylation level Y551, determined with a specific anti-pY551 antibody followed by secondary staining with FITC conjugated secondary antibody. The third column shows the cell periphery (plasma membrane) labeled with WGA-Alexa633, and the last column is an overlay. The quantification of colocalization between BTK and WGA by Pearson’s correlation coefficient (PCC) is shown in the bottom left of each panel. n= 22-25 over 3 independent experiments. scale bar = 30 μm. Boxplots represent quartiles. The data points outside the whisker range are set as outliers. The black line inside the box represents the median value. An unpaired two-tailed t-test was used to calculate significance. Boxplots were generated using Origin Pro 2020b. Image analysis was done using Fiji Ver 1.54f. The bottom right in each panel shows the schematic model of BTK activation by IP_6_.

### IP_6_ increases the cytosolic fraction of active BTK

We next focus on how the BTK^E41K^ activation profile at the membrane may be remodeled upon stimulating the transiently transfected CHO cells with IP_6_. We consider that the IP_6_ will enter the cell through the pinocytosis (43) and transphosphorylate Y551 of BTK^E41K^ by inducing an IP_6_-dependent dimerization (30). In Figures 4A-D, S3B, and S4, mCherry intensity indicates BTK expression level, FITC indicates the level of Y551 phosphorylation, and WGA-Alexa 633 indicates plasma membrane boundary. We observed that IP_6_ activates the cytosolic fraction of BTK wild-type (Figure 5A, 4A, and S4A). This is consistent with previous reports (22,33) that the wild-type BTK predominantly resides in the cytosol, which could be activated by IP_6_. In contrast, the addition of IP_6_ results in a significant redistribution of membrane-bound fraction of BTK^E41K^ to the cytosol (Figure 4B and S4B). The fraction of Y551 phosphorylation for the BTK^E41K^ mutant in the cytosol is significantly higher than the membrane-bound fraction (Figure 5A). The monomer control stays in the cytosol and cannot be activated by IP_6_ stimulation (Figure S3B-D).

**Figure 5:**
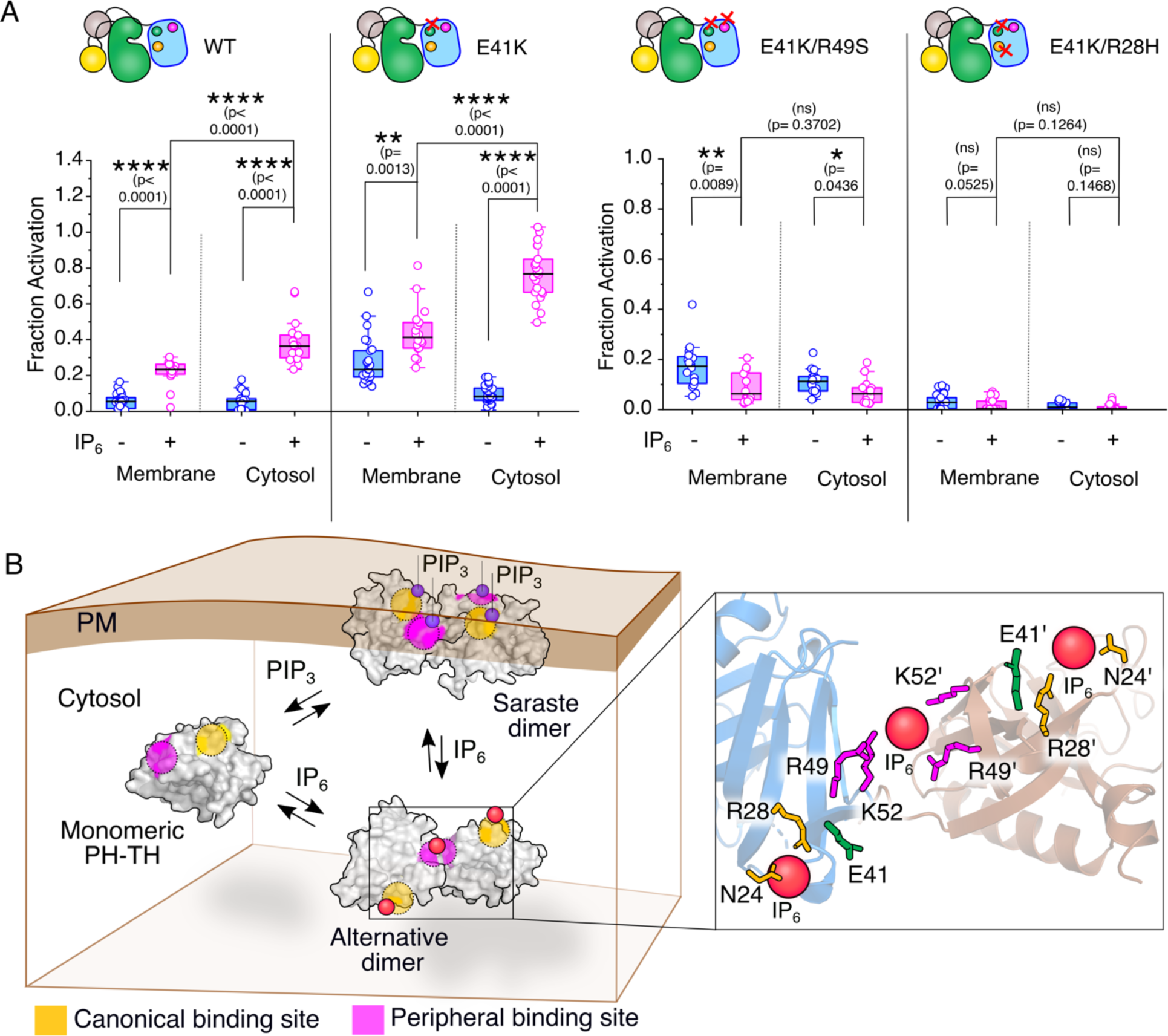
Determination of IP_6_-dependent autophosphorylation of BTK. A) Plots of fraction phosphorylated for the indicated BTK construct transiently expressed in CHO cell lines. The autophosphorylation level Y551 was determined with an anti-pY551 antibody. The phosphorylation of BTK localized at the plasma membrane and in the cytoplasm was measured in the presence and absence of IP_6_. The fraction was obtained by normalizing the pY551 level (FITC intensity) by the BTK expression level (mCherry intensity). n= 22-25 over 5 independent experiments. Boxplots represent quartiles. The data points outside the whisker range are set as outliers. The black line inside the box represents the median value. An unpaired two-tailed t-test was used to calculate significance. Boxplots were generated using Origin Pro 2020b. Image analysis was done using Fiji Ver 1.54f. B) The proposed model for the IP_6_-dependent dimerization of the PH-TH domain. In the absence of stimulation, the PH-TH domain remains in solution as a monomer. The binding of PIP_3_ at the canonical site recruits BTK to the membrane, and the Saraste dimer interface is stabilized by PIP_3_ binding to the peripheral site. The canonical and peripheral lipid binding sites are indicated with yellow and magenta, respectively. IP_6_ promotes an alternate dimer of the PH-TH domain shifting the equilibrium to a PH-TH dimer in solution. The residues in the canonical and peripheral binding sites are shown in the inset. The IP_6_ molecules are indicated by red balls.

Our *in-vitro* studies showed that the mutation at the peripheral lipid binding site (R49S) of PH-TH^E41K^ retains the ability to bind IP_6_ to the canonical site (Figure 3A and Table S1). We wonder if full-length BTK bearing the same mutation could be activated in cells. We observed that the BTK^E41K^ R49S mutant retained at the membrane even after being stimulated with IP_6_ and remains autoinhibited (Figure 4C and S4C). Our data suggests that the formation of the PH-TH Saraste dimer is crucial for BTK activation at the membrane or in the cytosol (30) (Figure 5B). However, the inability of IP_6_ to peel the R49S mutant of BTK^E41K^ from the plasma membrane indicates that the IP_6_ may not be competing with the PIP_3_ binding at the canonical lipid binding site. As anticipated, IP_6_ stimulation does not activate BTK^E41K^ mutated at the canonical binding site mutant (R28H) (Figure 4D and S3D). Suggesting that the two lipid-binding pockets of BTK are allosterically coupled to the Saraste dimer interface.

## Conclusions

BTK is indispensable for B-lymphocyte development and proliferation, particularly during the maturation of the pre-B-cell stage to the mature stage (7). The PH-TH domain of BTK functions as a master regulator. The activating mutation (E41K) (44,45) or loss of function mutant (XLA mutation) in the PH-TH domain leads to the manifestation of an immunodeficient phenotype (1,2). The β3-β4 segment of PH-TH plays an important role in the allosteric crosstalk during the membrane recruitment and dimerization of the kinase (29,30). One face of the β3-β4 segment constitutes the canonical lipid binding pocket for membrane localization (28). On the opposite side, the β3-β4 segment makes contact with an IP_6_ molecule (22) and laterally interacts with the neighboring PH-TH domains. Therefore, mutation (such as E41K) that stabilizes the β3-β4 segment (lower B-factor) (37) enhances the structural stability of the PH-TH domain (Figure 2E), thereby spontaneously activating the kinase.

Constitutive activation of BTK is often linked to several B-cell malignancies (7). Most of the BTK inhibitors are competitive inhibitors, competing for the ATP binding pocket. IP_6_ is the only BTK activator reported in the literature. IP_6_-mediated activation of BTK is distinct from IP_4,_ which works synergistically with the PIP_3_ binding for the full activation of Tec family protein tyrosine kinases Itk (46). In contrast, it was speculated that the increased cellular concentration of IP_6_ may outcompete the PIP_3_ and PH-TH binding at the membrane and shift the equilibrium more towards the cytosolic fraction (Figure 4B) (22,47). Counterintuitively, the IP_6_ only perturbs the membrane localization of the BTK^E41K^ but not the peripheral binding site mutant (R49S) (Figure 4C). Together, our data suggests that IP_6_ may not be competing with PIP_3_ at the membrane to bind the PH-TH domain.

The crystal structure of the PH-TH domain in the complex with IP_6_ (PDB ID: 4Y94) has four PH-TH domains present in the asymmetric unit (Figure S5) (22). Two PH-TH domains are dimerized by the lateral interaction between the Saraste dimer interface (Figure 2A) (28,37). The Saraste dimer is compatible with the simultaneous PIP_3_ binding to the canonical and the peripheral site at the membrane (29,30). An alternate homo dimer of PH-TH is also found (Figure S5) at the asymmetric unit of the PH-TH domain structure (PDB ID: 4Y94). The PH-TH domain dimerizes by an asymmetric interaction between the adjacent β3-β4 segments, forming a pocket. In the pocket formed at the interface of two PH-TH domains, one IP_6_ molecule is sandwiched, acting as a glue (Figure 5B). This dimer is only observed in the crystal structure of the IP_6_:PH-TH complex and not in the presence of other inositol polyphosphate. Such an IP_6_-dependent PH-TH dimer is incompatible with binding two PIP_3_ molecules simultaneously to the canonical lipid binding site at the membrane while retaining the IP_6_ molecule at the peripheral lipid binding site (Figure 5B). We speculate that IP_6_ may stabilize the alternate dimer over the Saraste dimer. Thus, shifting the equilibrium of the PH-TH dimer to the membrane-independent alternate conformation, thereby dislodging the BTK^E41K^ from the plasma membrane (Figure 4B and 5B). The peripheral site mutant (R49S) cannot form the alternate dimer and, hence, remains at the plasma membrane even in the presence of IP_6_ (Figure 4C).

Our data shows that the XLA mutation (R28H) impairs dimerization of the PH-TH domain and IP_6_-dependent phosphorylation of Y551 of BTK (Figure 4D and S3D). The R28H mutation may have impaired response to the T-cell-independent B-cell antigens, therefore, explaining why XLA patients are more susceptible to recurring bacterial and viral infection (1–4). We speculate that the ability of IP_6_ to activate BTK independent of BCR activation in our experiment opens a new opportunity to develop immune modulators.

### Experimental Procedures

#### Constructs

The PH-TH domain of BTK wildtype (amino acid residue number 4-170) cloned into pET28-SUMO vector was gifted from Prof. John Kuriyan, UC Berkeley. The pMSCV-BTK-mCherry construct was a gift from Hidde Ploegh (Addgene plasmid #50043). The full-length BTK was cloned into the pET28-SUMO vector. For the microscopy studies, the BTK-mCherry was cloned into a pCDNA vector. The point mutations in all the constructs were done by Site-directed mutagenesis.

#### Expression and purification of BTK PH-TH domain

The PH-TH domain of BTK cloned into pET28-SUMO vector was expressed in *E.coli* BL21-DE3 cells by inducing with 1 mM IPTG and at 18°C overnight (22). The cells were lysed in lysis buffer (25 mM Tris-Cl, 500 mM NaCl, 20 mM Imidazole, 5% Glycerol, 2 mM β-ME, pH 8.5). The protein was purified using a Ni-NTA column, and buffer exchanged to 25 mM Tris-Cl, 400 mM NaCl pH 8. His tag was removed by incubating with ULP1 protease for 12 hours, and the PH-TH domain was further purified by gel filtration chromatography. the protein was concentrated and stored at -80°C.

#### Expression and Purification of Full length BTK

The full length BTK cloned in pET28-SUMO vector was co-transformed with YopH and Trigger factor in *E.coli* BL21-DE3 cells (22). Briefly, the cells were grown in LB supplemented with 50 mg/ml Kanamycin, 35 mg/ml Chloramphenicol, and 50 mg/ml. The BTK expression was induced with 1 mM IPTG and was grown at 18°C for 16 hrs. The cells were lysed in lysis buffer (25 mM Tris-Cl, 500 mM NaCl, 20 mM Imidazole, 5% Glycerol, 2 mM β-ME, pH 8.5) and the protein was purified using Ni-NTA column. His tag was removed by digesting BTK with ULP1 protease for 12 hours. The undigested protein was then removed by passing through the second Ni-NTA column and flow through was concentrated and stored at -80°C.

#### Isothermal Titration Calorimetry

The isothermal titration calorimetry experiments were performed using MicroCal ITC (GE Healthcare) and MicroCal PEAQ-ITC (Malvern). The PH-TH domain was buffer exchanged to 20 mM HEPES, 150 mM NaCl, pH 7.5. During the titration, 25 µM protein in the cell was titrated with a ligand at the protein-to-ligand ratio of 1: 5.5. All ITC titrations were performed at 20°C. The initial injection of 0.5 µl was excluded from the data analysis. During the titration, 3 µl of ligand was injected in 13, and each injection step was separated by 180 secs. The protein solution was stirred at 300 rpm during the titration.

#### Gel Filtration Assay

Gel filtration assay of various constructs of the PH-TH domain was performed using the Superdex-200 10/300 GL column. For the apo PH-TH domain, the column was equilibrated with running buffer (25 mM Tris-Cl, 100 mM NaCl, 1 mM DTT, pH 7.2), and 500 µl of 75 µM protein was loaded into the column, at a flow rate of 0.3 ml/min and at 8°C. The elution profile was recorded, and the retention time of the protein was calculated. For the ligand-bound samples, the protein was mixed with ligand at a protein-to-ligand ratio of 1:4, and incubated at 4°C for 15 minutes before loading into the gel filtration column. Before loading the sample, the column was pre-equilibrated with a running buffer containing the ligand.

#### Dynamic Light Scattering

The Dynamic Light Scattering experiments were conducted using Malvern Zetasizer DLS Detectors using 1.5 mL plastic cuvettes. The samples were prepared in a buffer containing 25 mM Tris-Cl, 100 mM NaCl, 1 mM DTT, and pH 7.2. The experiment included 3 measurements of 10 runs each. The theoretical calculations of the hydrodynamic radius of the proteins were performed using Hydropro (48).

#### Thermal denaturation by Circular Dichroism (CD) Spectroscopy

The melting temperature of the PH-TH domain was determined from the CD spectrum recorded at increasing temperatures (20° to 76°C). For each sample, the CD spectra from 300 to 190 nm were recorded using a Jasco-J1500 spectropolarimeter with a temperature increment of 4°C. CD spectra were recorded with 5μM protein in 20 mM Sodium phosphate buffer pH 7.4 and in the absence or in the presence of an equimolar ligand. The measured ellipticity data was converted into molar ellipticity (49).

#### Autophosphorylation assay by immunoblot

The activation of full-length BTK in response to the ligand was determined from the autophosphorylation Y551 or total phosphotyrosine level. Before each reaction, 2 µM BTK was incubated with 100 µM ligand for 10 minutes. The phosphorylation reaction was performed at 25°C in a buffer containing 25 mM Tris-Cl (pH 7.5), 2 mM ATP, 10 mM MgCl_2,_ and 1 mM Na_3_VO_4_. The reaction was quenched with SDS–PAGE loading buffer. The autophosphorylation levels of full-length BTK were detected by immunoblot (22). The amount of BTK loaded was determined with anti-BTK antibody. Densitometric analysis of immunoblots was performed using Fiji software (50).

#### Cell-based assay

The IP_6_-mediated activation of BTK was determined by immunofluorescence and immunoblot in Chinese Hamster Ovary (CHO) cell line transiently transfected with BTK-mCherry cloned in pCDNA. Before activation, transfected cells were serum-starved for six hours in opti-MEM, followed by incubation with 500µM IP_6_ for 1 hour in opti-MEM. For Immunoblot, cells were washed with ice-cold PBS and then incubated in RIPA buffer [10 mM Tris-Cl (pH 8.0), 140 mM NaCl, 1 mM EDTA, 0.1% SDS, and 1% Triton X-100] containing protease and phosphatase inhibitors (2 mM Benzamidine, 1 mM PMSF and 1 mM Sodium orthovanadate) for 10 mins. The cell lysates were sonicated, and protein samples were prepared by heating with 5X loading buffer and resolved in a 6% SDS-PAGE. Protein samples were transferred onto the PVDF membrane at 15 V for 1 hour, followed by blocking with 5% skimmed milk in 1X TBS containing 0.1% TWEEN-20 for 1 hr at room temperature. After blocking, the blot was incubated overnight at 4°C with the primary antibody (1:1000) diluted in 3% skimmed milk. After incubation, the blot was washed three times with 1X TBST (0.1% TWEEN-20), followed by incubation with a secondary antibody (1:2000) diluted in 3% skimmed milk. Blots were washed three times with 1X TBST and developed using the ClarityTM Western ECL substrate kit (Bio-Rad). Densitometry analysis of immunoblots was performed using Fiji Ver 1.54f (50).

For the immunofluorescence study, Cells were washed with PBS once, followed by incubation with WGA (20ug/ml) in PBS at 37°C for 1 min. After incubation, cells were washed with PBS and immediately fixed with 4% paraformaldehyde in PBS for 30 minutes at room temperature. After fixation, cells were washed five times with 1x PBS and permeabilized with 0.2% PBST for 5 min at room temperature. Cells were blocked with 1% BSA for 1 hour at room temperature, followed by staining with anti-551pY primary antibody (1:100 dilution) in blocking buffer overnight at 4⁰C. After incubation, cells were incubated with a FITC-conjugated secondary antibody in blocking buffer for 2 hours at room temperature. Finally, coverslips were mounted with prolonged gold. Coverslips were washed three times with 1X PBS in between each step. Details of the antibody used are summarized in Table S3.

##### Microscopy and image analysis

All images were acquired with the Leica SP8 confocal platform using an oil immersion HC PL APO CS2 63× objective (NA 1.4) at 3.0X digital zoom. Image scanning was done in bidirectional mode at 500 Hz. The level of BTK on the membrane or cytoplasm was determined from the intensity of mCherry measured at λ_ex_= 552 nm and λ_em_= 576-651 nm, respectively. The Y551 phosphorylation was determined from the intensity of FITC tagged to the secondary antibody and measured at λ_ex_= 488 nm and λ_em_= 505-531 nm. The plasma membrane was stained with Wheat Germ Agglutinin (WGA) fused to Alexa 633 and imaged at λ_ex_= 633 nm, λ_em_= 647-692 nm, respectively. ROIs were drawn manually using the Fiji freehand tool (Ver 1.54f) (50). ROI1 marked the outer boundary of the cell perimeter labeled with WGA, ROI2 marked the inner boundary of the cell perimeter, and ROI3 marked the nucleus. The ROI1 provides the total cell intensity. The intensity of mCherry or FITC at the membrane was determined by subtracting the respective intensities of ROI2 from ROI1. Whereas cytoplasmic intensity of mCherry or FITC was determined by subtracting the respective intensities of ROI3 from ROI2. The membrane localization of the BTK-mCherry construct was determined from the quantification of Pearson’s colocalization coefficient (PCC) (42).

## Supporting Information

The Supporting Information contains Figures S1-S5 and Tables S1-S3.

## Acknowledgements

The authors thank Prof. Arnab Gupta for access to the confocal microscope and Prosad Kumar Das for helping with the expression of full-length BTK-mCherry in the initial stage of the experiment.

## Author Contributions

The manuscript was written through the contribution of all authors. All authors have approved the final version of the manuscript. RD, SC, and MPC designed the experiments. SC, SR, KG, and MPC performed the Biochemical experiments and cell-based studies. RD, SC, SR, KG, and MPC wrote the manuscript.

## Funding and additional information

The authors thank research funding from IISER Kolkata, infrastructural facilities supported by IISER Kolkata, and DST-FIST (SR/FST/LS-II/2017/93(c)). This work is supported by grants from SERB (ECR/2015/000142) and (CRG/2020/000437).

## Data Availability Statement

All the relevant data are contained within this article and in the supporting information.

## References

1. Tsukada, S., Saffran, D. C., Rawlings, D. J., Parolini, O., Allen, R. C., Klisak, I., Sparkes, R. S., Kubagawa, H., Mohandas, T., Quan, S., and, et al. (1993) Deficient expression of a B cell cytoplasmic tyrosine kinase in human X-linked agammaglobulinemia. Cell 72, 279–290

2. Vetrie, D., Vorechovsky, I., Sideras, P., Holland, J., Davies, A., Flinter, F., Hammarstrom, L., Kinnon, C., Levinsky, R., Bobrow, M., and, et al. (1993) The gene involved in X-linked agammaglobulinaemia is a member of the src family of protein-tyrosine kinases. Nature 361, 226–233

3. Thomas, J. D., Sideras, P., Smith, C. I., Vorechovsky, I., Chapman, V., and Paul, W. E. (1993) Colocalization of X-linked agammaglobulinemia and X-linked immunodeficiency genes. Science 261, 355–358

4. Fukuda, M., Kojima, T., Kabayama, H., and Mikoshiba, K. (1996) Mutation of the pleckstrin homology domain of Bruton’s tyrosine kinase in immunodeficiency impaired inositol 1, 3, 4, 5-tetrakisphosphate binding capacity. Journal of Biological Chemistry 271, 30303–30306

5. Woyach, J. A., Bojnik, E., Ruppert, A. S., Stefanovski, M. R., Goettl, V. M., Smucker, K. A., Smith, L. L., Dubovsky, J. A., Towns, W. H., MacMurray, J., Harrington, B. K., Davis, M. E., Gobessi, S., Laurenti, L., Chang, B. Y., Buggy, J. J., Efremov, D. G., Byrd, J. C., and Johnson, A. J. (2014) Bruton’s tyrosine kinase (BTK) function is important to the development and expansion of chronic lymphocytic leukemia (CLL). Blood 123, 1207–1213

6. Davis, R. E., Ngo, V. N., Lenz, G., Tolar, P., Young, R. M., Romesser, P. B., Kohlhammer, H., Lamy, L., Zhao, H., Yang, Y., Xu, W., Shaffer, A. L., Wright, G., Xiao, W., Powell, J., Jiang, J. K., Thomas, C. J., Rosenwald, A., Ott, G., Muller-Hermelink, H. K., Gascoyne, R. D., Connors, J. M., Johnson, N. A., Rimsza, L. M., Campo, E., Jaffe, E. S., Wilson, W. H., Delabie, J., Smeland, E. B., Fisher, R. I., Braziel, R. M., Tubbs, R. R., Cook, J. R., Weisenburger, D. D., Chan, W. C., Pierce, S. K., and Staudt, L. M. (2010) Chronic active B-cell-receptor signalling in diffuse large B-cell lymphoma. Nature 463, 88–92

7. Hendriks, R. W., Yuvaraj, S., and Kil, L. P. (2014) Targeting Bruton’s tyrosine kinase in B cell malignancies. Nat Rev Cancer 14, 219–232

8. Joseph, R. E., Amatya, N., Fulton, D. B., Engen, J. R., Wales, T. E., and Andreotti, A. (2020) Differential impact of BTK active site inhibitors on the conformational state of full-length BTK. Elife 9

9. Byrd, J. C., Furman, R. R., Coutre, S. E., Flinn, I. W., Burger, J. A., Blum, K. A., Grant, B., Sharman, J. P., Coleman, M., Wierda, W. G., Jones, J. A., Zhao, W., Heerema, N. A., Johnson, A. J., Sukbuntherng, J., Chang, B. Y., Clow, F., Hedrick, E., Buggy, J. J., James, D. F., and O’Brien, S. (2013) Targeting BTK with ibrutinib in relapsed chronic lymphocytic leukemia. N Engl J Med 369, 32–42

10. Wang, M. L., Rule, S., Martin, P., Goy, A., Auer, R., Kahl, B. S., Jurczak, W., Advani, R. H., Romaguera, J. E., Williams, M. E., Barrientos, J. C., Chmielowska, E., Radford, J., Stilgenbauer, S., Dreyling, M., Jedrzejczak, W. W., Johnson, P., Spurgeon, S. E., Li, L., Zhang, L., Newberry, K., Ou, Z., Cheng, N., Fang, B., McGreivy, J., Clow, F., Buggy, J. J., Chang, B. Y., Beaupre, D. M., Kunkel, L. A., and Blum, K. A. (2013) Targeting BTK with ibrutinib in relapsed or refractory mantle-cell lymphoma. N Engl J Med 369, 507–516

11. Lee, K. G., Xu, S., Kang, Z. H., Huo, J., Huang, M., Liu, D., Takeuchi, O., Akira, S., and Lam, K. P. (2012) Bruton’s tyrosine kinase phosphorylates Toll-like receptor 3 to initiate antiviral response. Proc Natl Acad Sci U S A 109, 5791–5796

12. Desiderio, S. (1997) Role of Btk in B cell development and signaling. Curr Opin Immunol 9, 534–540

13. Wahl, M. I., Fluckiger, A. C., Kato, R. M., Park, H., Witte, O. N., and Rawlings, D. J. (1997) Phosphorylation of two regulatory tyrosine residues in the activation of Bruton’s tyrosine kinase via alternative receptors. Proc Natl Acad Sci U S A 94, 11526–11533

14. Salim, K., Bottomley, M. J., Querfurth, E., Zvelebil, M., Gout, I., Scaife, R., Margolis, R., Gigg, R., Smith, C., and Driscoll, P. (1996) Distinct specificity in the recognition of phosphoinositides by the pleckstrin homology domains of dynamin and Bruton’s tyrosine kinase. The EMBO journal 15, 6241–6250

15. Rawlings, D. J., and Witte, O. N. (1995) The Btk subfamily of cytoplasmic tyrosine kinases: structure, regulation and function. Semin Immunol 7, 237–246

16. Nagar, B., Hantschel, O., Young, M. A., Scheffzek, K., Veach, D., Bornmann, W., Clarkson, B., Superti-Furga, G., and Kuriyan, J. (2003) Structural basis for the autoinhibition of c-Abl tyrosine kinase. Cell 112, 859–871

17. Xu, W., Harrison, S. C., and Eck, M. J. (1997) Three-dimensional structure of the tyrosine kinase c-Src. Nature 385, 595–602

18. Sicheri, F., Moarefi, I., and Kuriyan, J. (1997) Crystal structure of the Src family tyrosine kinase Hck. Nature 385, 602–609

19. Sideras, P., and Smith, C. I. (1995) Molecular and cellular aspects of X-linked agammaglobulinemia. Adv Immunol 59, 135–223

20. Mohamed, A. J., Yu, L., Backesjo, C. M., Vargas, L., Faryal, R., Aints, A., Christensson, B., Berglof, A., Vihinen, M., Nore, B. F., and Smith, C. I. (2009) Bruton’s tyrosine kinase (Btk): function, regulation, and transformation with special emphasis on the PH domain. Immunol Rev 228, 58–73

21. Joseph, R. E., Min, L., and Andreotti, A. H. (2007) The linker between SH2 and kinase domains positively regulates catalysis of the Tec family kinases. Biochemistry 46, 5455–5462

22. Wang, Q., Vogan, E. M., Nocka, L. M., Rosen, C. E., Zorn, J. A., Harrison, S. C., and Kuriyan, J. (2015) Autoinhibition of Bruton’s tyrosine kinase (Btk) and activation by soluble inositol hexakisphosphate. Elife 4, e06074

23. Devkota, S., Joseph, R. E., Boyken, S. E., Fulton, D. B., and Andreotti, A. H. (2017) An Autoinhibitory Role for the Pleckstrin Homology Domain of Interleukin-2-Inducible Tyrosine Kinase and Its Interplay with Canonical Phospholipid Recognition. Biochemistry 56, 2938–2949

24. Amatya, N., Wales, T. E., Kwon, A., Yeung, W., Joseph, R. E., Fulton, D. B., Kannan, N., Engen, J. R., and Andreotti, A. H. (2019) Lipid-targeting pleckstrin homology domain turns its autoinhibitory face toward the TEC kinases. Proceedings of the National Academy of Sciences 116, 21539–21544

25. Joseph, R. E., Wales, T. E., Fulton, D. B., Engen, J. R., and Andreotti, A. H. (2017) Achieving a Graded Immune Response: BTK Adopts a Range of Active/Inactive Conformations Dictated by Multiple Interdomain Contacts. Structure 25, 1481–1494 e1484

26. Aoki, Y., Isselbacher, K. J., and Pillai, S. (1994) Bruton tyrosine kinase is tyrosine phosphorylated and activated in pre-B lymphocytes and receptor-ligated B cells. Proc Natl Acad Sci U S A 91, 10606–10609

27. de Weers, M., Brouns, G. S., Hinshelwood, S., Kinnon, C., Schuurman, R. K., Hendriks, R. W., and Borst, J. (1994) B-cell antigen receptor stimulation activates the human Bruton’s tyrosine kinase, which is deficient in X-linked agammaglobulinemia. J Biol Chem 269, 23857–23860

28. Hyvonen, M., and Saraste, M. (1997) Structure of the PH domain and Btk motif from Bruton’s tyrosine kinase: molecular explanations for X-linked agammaglobulinaemia. EMBO J 16, 3396–3404

29. Wang, Q., Pechersky, Y., Sagawa, S., Pan, A. C., and Shaw, D. E. (2019) Structural mechanism for Bruton’s tyrosine kinase activation at the cell membrane. Proceedings of the National Academy of Sciences 116, 9390–9399

30. Chung, J. K., Nocka, L. M., Decker, A., Wang, Q., Kadlecek, T. A., Weiss, A., Kuriyan, J., and Groves, J. T. (2019) Switch-like activation of Bruton’s tyrosine kinase by membrane-mediated dimerization. Proc Natl Acad Sci U S A 116, 10798–10803

31. Park, H., Wahl, M. I., Afar, D. E., Turck, C. W., Rawlings, D. J., Tam, C., Scharenberg, A. M., Kinet, J.-P., and Witte, O. N. (1996) Regulation of Btk function by a major autophosphorylation site within the SH3 domain. Immunity 4, 515–525

32. Rawlings, D. J., Scharenberg, A. M., Park, H., Wahl, M. I., Lin, S., Kato, R. M., Fluckiger, A.-C., Witte, O. N., and Kinet, J.-P. (1996) Activation of BTK by a phosphorylation mechanism initiated by SRC family kinases. Science 271, 822–825

33. Kim, W., Kim, E., Min, H., Kim, M. G., Eisenbeis, V. B., Dutta, A. K., Pavlovic, I., Jessen, H. J., Kim, S., and Seong, R. H. (2019) Inositol polyphosphates promote T cell-independent humoral immunity via the regulation of Bruton’s tyrosine kinase. Proceedings of the National Academy of Sciences 116, 12952–12957

34. Saiardi, A., Erdjument-Bromage, H., Snowman, A. M., Tempst, P., and Snyder, S. H. (1999) Synthesis of diphosphoinositol pentakisphosphate by a newly identified family of higher inositol polyphosphate kinases. Curr Biol 9, 1323–1326

35. Li, T., Tsukada, S., Satterthwaite, A., Havlik, M. H., Park, H., Takatsu, K., and Witte, O. N. (1995) Activation of Bruton’s tyrosine kinase (BTK) by a point mutation in its pleckstrin homology (PH) domain. Immunity 2, 451–460

36. Kanegane, H., Futatani, T., Wang, Y., Nomura, K., Shinozaki, K., Matsukura, H., Kubota, T., Tsukada, S., and Miyawaki, T. (2001) Clinical and mutational characteristics of X-linked agammaglobulinemia and its carrier identified by flow cytometric assessment combined with genetic analysis. J Allergy Clin Immunol 108, 1012–1020

37. Baraldi, E., Carugo, K. D., Hyvönen, M., Surdo, P. L., Riley, A. M., Potter, B. V., O’Brien, R., Ladbury, J. E., and Saraste, M. (1999) Structure of the PH domain from Bruton’s tyrosine kinase in complex with inositol 1, 3, 4, 5-tetrakisphosphate. Structure 7, 449–460

38. Várnai, P., Rother, K. I., and Balla, T. (1999) Phosphatidylinositol 3-kinase-dependent membrane association of the Bruton’s tyrosine kinase pleckstrin homology domain visualized in single living cells. Journal of Biological Chemistry 274, 10983–10989

39. de Weers, M., Mensink, R. G., Kraakman, M. E., Schuurman, R. K., and Hendriks, R. W. (1994) Mutation analysis of the Bruton’s tyrosine kinase gene in X-linked agammaglobulinemia: identification of a mutation which affects the same codon as is altered in immunodeficient xid mice. Hum Mol Genet 3, 161–166

40. Saouaf, S. J., Mahajan, S., Rowley, R. B., Kut, S. A., Fargnoli, J., Burkhardt, A. L., Tsukada, S., Witte, O. N., and Bolen, J. B. (1994) Temporal differences in the activation of three classes of non-transmembrane protein tyrosine kinases following B-cell antigen receptor surface engagement. Proc Natl Acad Sci U S A 91, 9524–9528

41. Chazotte, B. (2011) Labeling membrane glycoproteins or glycolipids with fluorescent wheat germ agglutinin. Cold Spring Harbor Protocols 2011, pdb. prot5623

42. Kar, S., Sen, S., Maji, S., Saraf, D., Ruturaj, Paul, R., Dutt, S., Mondal, B., Rodriguez-Boulan, E., Schreiner, R., Sengupta, D., and Gupta, A. (2022) Copper(II) import and reduction are dependent on His-Met clusters in the extracellular amino terminus of human copper transporter-1. J Biol Chem 298, 101631

43. Vucenik, I., and Shamsuddin, A. M. (1994) [3H]inositol hexaphosphate (phytic acid) is rapidly absorbed and metabolized by murine and human malignant cells in vitro. J Nutr 124, 861–868

44. Maas, A., Dingjan, G. M., Grosveld, F., and Hendriks, R. W. (1999) Early arrest in B cell development in transgenic mice that express the E41K Bruton’s tyrosine kinase mutant under the control of the CD19 promoter region. J Immunol 162, 6526–6533

45. Dingjan, G. M., Maas, A., Nawijn, M. C., Smit, L., Voerman, J. S., Grosveld, F., and Hendriks, R. W. (1998) Severe B cell deficiency and disrupted splenic architecture in transgenic mice expressing the E41K mutated form of Bruton’s tyrosine kinase. EMBO J 17, 5309–5320

46. Huang, Y. H., Grasis, J. A., Miller, A. T., Xu, R., Soonthornvacharin, S., Andreotti, A. H., Tsoukas, C. D., Cooke, M. P., and Sauer, K. (2007) Positive regulation of Itk PH domain function by soluble IP4. Science 316, 886–889

47. Chakraborty, A., Koldobskiy, M. A., Bello, N. T., Maxwell, M., Potter, J. J., Juluri, K. R., Maag, D., Kim, S., Huang, A. S., Dailey, M. J., Saleh, M., Snowman, A. M., Moran, T. H., Mezey, E., and Snyder, S. H. (2010) Inositol pyrophosphates inhibit Akt signaling, thereby regulating insulin sensitivity and weight gain. Cell 143, 897–910

48. Ortega, A., Amorós, D., and De La Torre, J. G. (2011) Prediction of hydrodynamic and other solution properties of rigid proteins from atomic-and residue-level models. Biophysical journal 101, 892–898

49. Greenfield, N. J. (2006) Using circular dichroism collected as a function of temperature to determine the thermodynamics of protein unfolding and binding interactions. Nat Protoc 1, 2527–2535

50. Schindelin, J., Arganda-Carreras, I., Frise, E., Kaynig, V., Longair, M., Pietzsch, T., Preibisch, S., Rueden, C., Saalfeld, S., Schmid, B., Tinevez, J. Y., White, D. J., Hartenstein, V., Eliceiri, K., Tomancak, P., and Cardona, A. (2012) Fiji: an open-source platform for biological-image analysis. Nat Methods 9, 676–682

