## Supplemental Information for "E41K Mutation Activates Bruton’s Tyrosine Kinase by Stabilizing an Inositol Hexakisphosphate dependent Invisible Dimer"

**Manuscript**

Corresponding authors

Rahul Das:

**Keywords:** Kinase, Cell signaling, BTK, Inositol Hexakisphosphate, and B-cell receptor

**Table S1: Thermodynamic parameters for binding of $IP_6$ to the BTK PH-TH variants derived from Isothermal Titration Calorimetry**

| PH-TH Constructs | $K_{d1}$ (nM) | $\Delta H_1$ (Cal/mole) | $K_{d2}$ (nM) | $\Delta H_2$ (Cal/mole) |
|---|---|---|---|---|
| E41K | 152 ± 18 | - 4749 ± 159 | 1275 ± 366 | -5098 ± 2360 |
| E41K/R28H | 2421 ± 151 | - 1562 ± 1008 | - | - |
| E41K/R49S | 622 ± 77 | - 4160 ± 550 | - | - |
| E41K/R28H/R49S | - | - | - | - |
| R28H/R49S | - | - | - | - |

**Table S2: $\Delta G_{unfolding}$ and $\Delta\Delta G_{unfolding}$ of Apo and $IP_6$ bound PH-TH domain of BTK**

| PH-TH Constructs | $\Delta G_{unfolding}$ (Cal/mole) | $\Delta\Delta G_{unfolding}$ (Cal/mole) |
|---|---|---|
| WT Apo | 341.670 ± 504 | - |
| WT + $IP_6$ | -686.971 ± 188 | -1028.642 ± 153.88 |
| E41K Apo | 469.713 ± 196 | 128.042 ± 196.06 |
| E41K + $IP_6$ | -1307.023 ± 954 | -1648.694 ± 779.38 |
| E41K + $IP_3$ | 326.697 ± 145 | -14.972 ± 84.05 |
| E41K/R28H/R49S Apo | 2264.570 ± 366 | 1922.9 ± 299.6 |
| E41K/R28H/R49S + $IP_6$ | 2231.412 ± 553 | 1889.741 ± 451.66 |

**Table S3: Details of antibodies**

| Antibodies | Source | |
|---|---|---|
| BTK (D3H5) Rabbit mAb | Cell Signaling Technology, Danvers, MA, USA | Cat # 8547S Lot: 13 |
| Recombinant Anti-BTK (phospho Y551) antibody [EP267Y] | Abcam (Waltham, MA 02453, USA) | Cat # ab40770 GR111548-5 |
| Recombinant Anti-Phosphotyrosine antibody [EPR16871] | Abcam (Waltham, MA 02453, USA) | Cat # ab179530 GR198792-25 |
| Rabbit HRP Secondary antibody | Abcam (Waltham, MA 02453, USA) | Cat # 50095 Lot: 2960660 |
| GAPDH rabbit polyclonal antibody | BioBharati Life Science Pvt Ltd | Cat # BB-AB0060 Lot: 011501 |
| Rabbit FITC-conjugated secondary antibody | Abcam (Waltham, MA 02453, USA) | Cat # Ab6885 Lot: GR3391568-I |

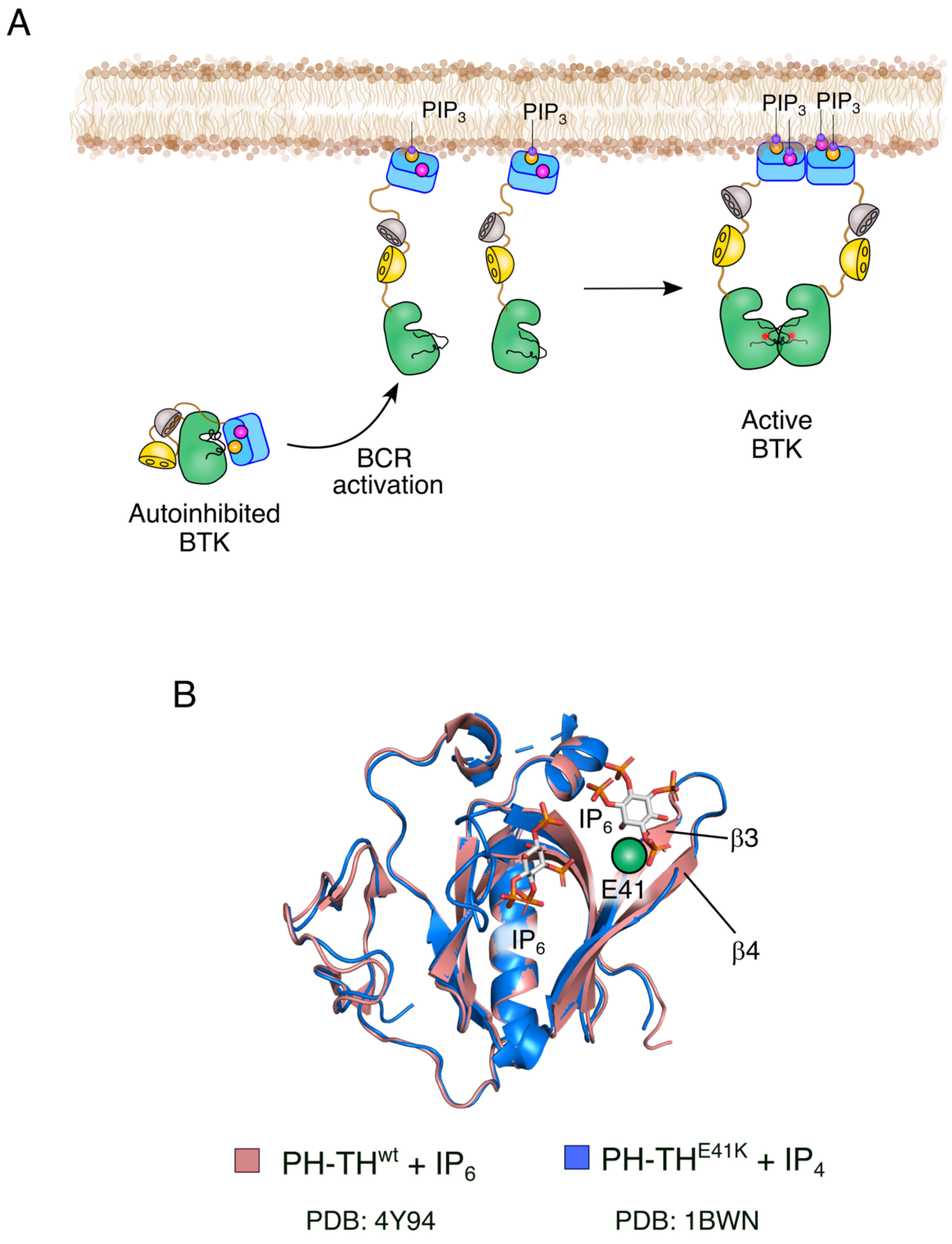


**Figure S1. BCR-mediated activation of BTK**

(A) Schematic representation of $PIP_3$ mediated membrane recruitment of BTK upon BCR activation.

(B) Structural alignment of PH-TH$^{WT}$ in complex with $IP_6$ (PDB ID: 4Y94) (1) and PH-TH$^{E41K}$ domain in complex with $IP_4$ (PDB ID: 1BWN) (2).

A

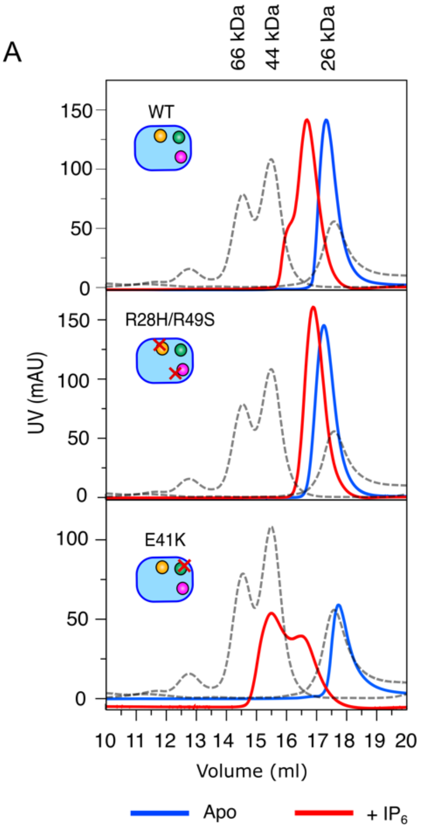


B

| Oligomeric state of PH-TH | Molecular Weight (kD) | Expected Hydrodynamic radius(nm) |
|---|---|---|
| Monomer | 19 | 2.20 |
| Dimer | 38 | 3.00 |
| Tetramer | 76 | 3.76 |

C

| PH-TH | Hydrodynamic radius(nm) - $IP_6$ | Hydrodynamic radius(nm) + $IP_6$ |
|---|---|---|
| WT | 2.42 ± 0.46 | 2.18 ± 0.10 |
| E41K | 2.57 ± 0.67 | 3.14 ± 0.48 |
| R28H/R49S | 2.11 ± 0.30 | 2.20 ± 0.19 |
| E41K/R28H/R49S | 2.55 ± 0.21 | 2.67 ± 0.27 |

D

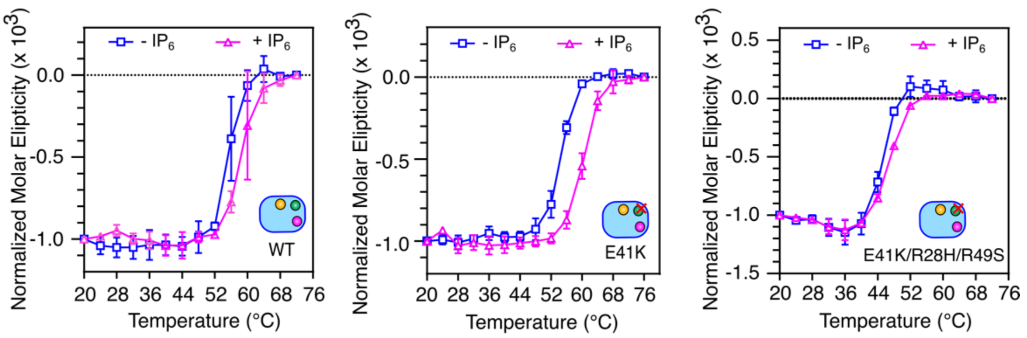


**Figure S2. Characterization the $IP_6$ dependent PH-TH dimer in solution**

A) Representative gel-filtration elution profiles of indicated PH-TH constructs measured in the presence or absence of $IP_6$. The dotted line represents the elution profile of a standard protein mixture comprised of BSA (66 kDa), Ovalbumin (44 kDa), and ULP1 (26 kDa). The plots were generated by XMGRACE Ver 5.1.25.

B) Predicted Hydrodynamic radius of different oligomeric states of PH-TH domain of BTK calculations using Hydropro software (3).

(C) The hydrodynamic radius of BTK PH-TH domain variants in the presence or absence of $IP_6$ was measured using Dynamic Light Scattering (DLS).

(D) Thermal denaturation profiles of indicated PH-TH constructs in the presence (magenta) or absence (cyan) of $IP_6$. Data are presented as mean values ± SD from three independent experiments. The solid line represents the fitting to the Boltzmann Sigmoidal equation using GraphPad Prism version 9.5.1(GraphPad Software, LLC.). Data analyses were performed using GraphPad Prism version 9.5.1.

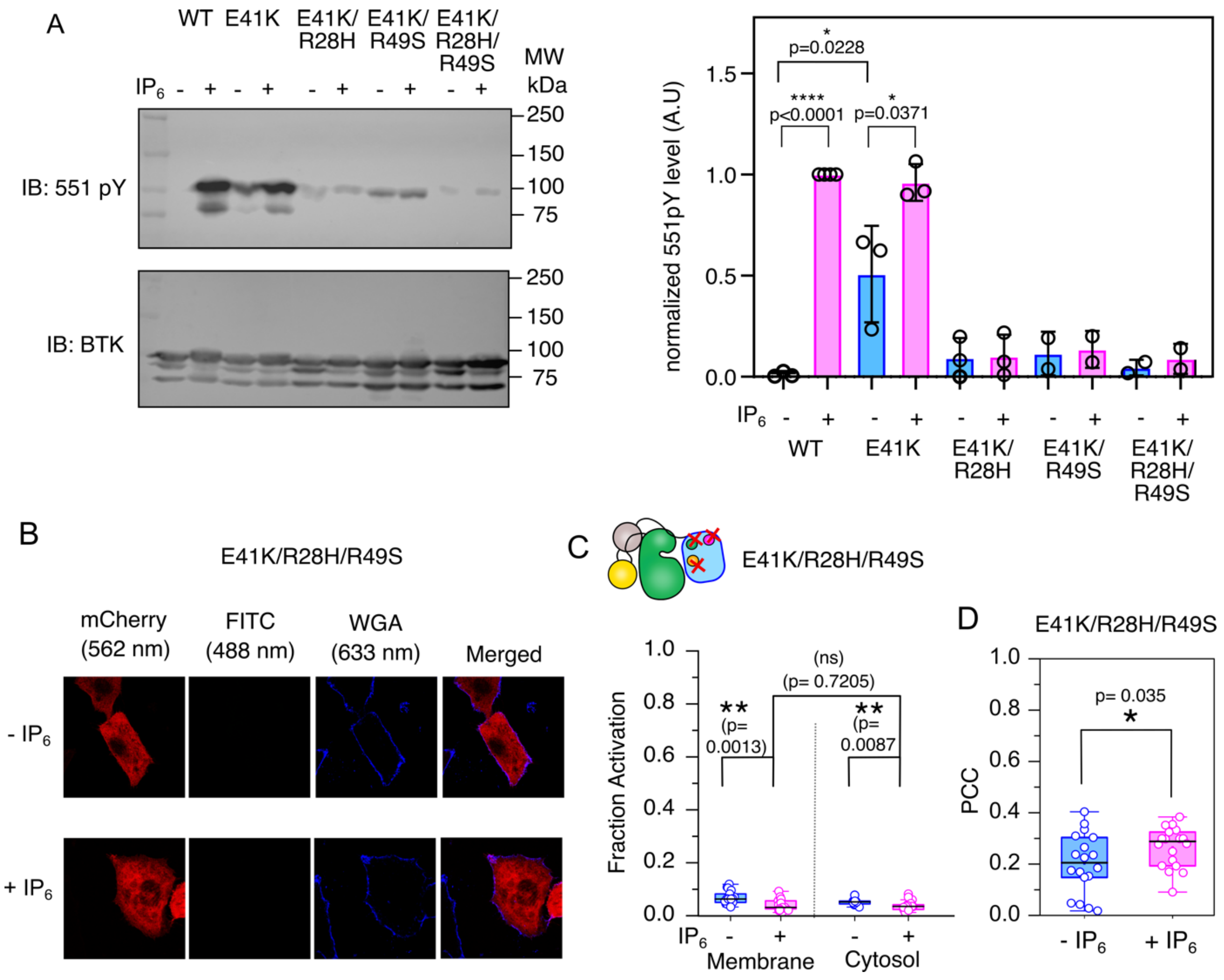


**Figure S3. $IP_6$-mediated activation of BTK in CHO cells**

(A) The left panel is the representative immunoblot of Y551 phosphorylation level in the indicated construct of BTK transiently expressed in the CHO cell line in the presence and absence of $IP_6$. The densitometric analysis of the immunoblot is on the right. Data are presented as mean values ± SD from five independent experiments. Data analyses were performed using GraphPad Prism version 9.5.1.

B) Confocal images of BTK E41K/R28H/R49S mutant in the presence (bottom) or absence (up) of $IP_6$. The BTK expression level is shown in red ($\lambda_{ex}$= 552 nm, $\lambda_{em}$= 586-651 nm), and the phosphorylation status is shown in green ($\lambda_{ex}$= 488 nm, $\lambda_{em}$= 505-531 nm). The blue represents the plasma membrane stained with Wheat Germ Agglutinin (WGA) fused to Alexa 633 ($\lambda_{ex}$= 633 nm, $\lambda_{em}$= 647-692 nm).

C) Quantification of colocalization of BTK E41K/R28H/R49S mutant and WGA by Pearson's correlation coefficient. n= 22-25 over 3 independent experiments. Boxplots represent quartiles. The data points outside the whisker range are set as outliers. The black line inside the box represents the median value. boxplots were generated using Origin Pro 2020b. Image analysis was done using Fiji Ver 1.54f (4).

D) The plot of fraction phosphorylated for the BTK E41K/R28H/R49S mutant transiently expressed in CHO cell lines. n= 22-25 over 5 independent experiments. Boxplots represent quartiles. The data points outside the whisker range are set as outliers. The black line inside the box represents the median value. boxplots were generated using Origin Pro 2020b. An unpaired two-tailed t-test was used to calculate significance.

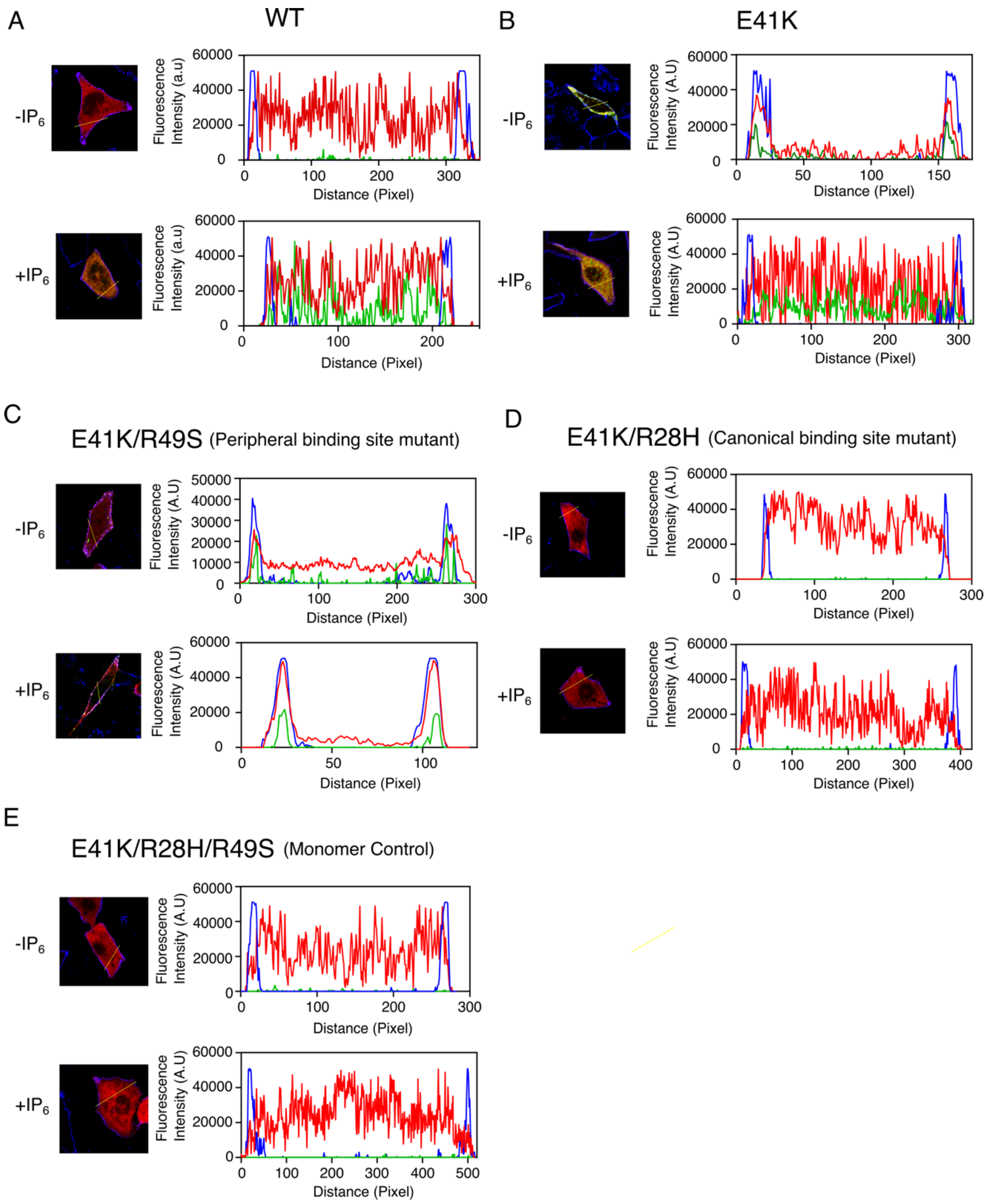


**Figure S4. (A-E)** Intensity plots of indicated BTK constructs transiently expressed in CHO cell line, in the presence or absence of $IP_6$. The BTK expression level is shown in red, the phosphorylation level of Y551 is shown in. The blue represents the plasma membrane stained with Wheat Germ Agglutinin (WGA) fused to Alexa 633. Image analysis was done using Fiji Ver 1.54f. The plots were generated using GraphPad Prism version 9.5.1.

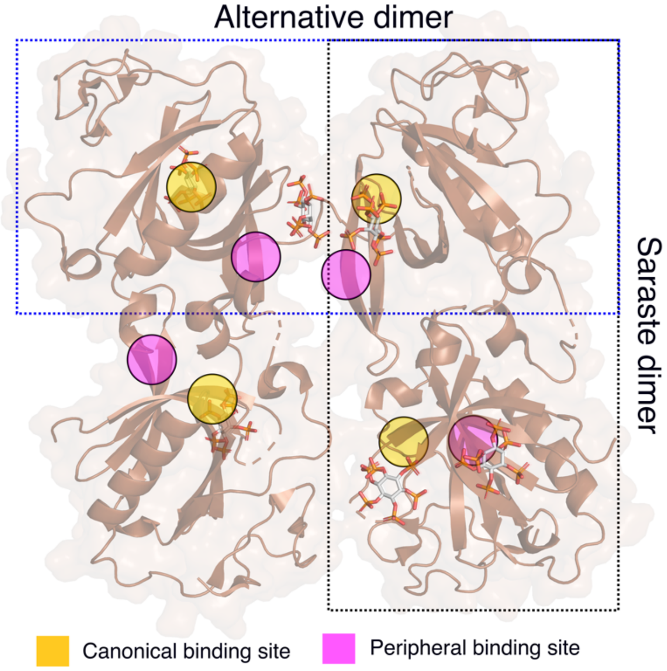


**Figure S5.** Cartoon representation of PH-TH molecule in the asymmetric unit of the PH-TH: $IP_6$ crystal structure (PDB: 4Y94) (1). The Saraste dimer and the alternative dimers are shown in the box.

### Supplementary References


1. Wang, Q., Vogan, E. M., Nocka, L. M., Rosen, C. E., Zorn, J. A., Harrison, S. C., and Kuriyan, J. (2015) Autoinhibition of Bruton's tyrosine kinase (Btk) and activation by soluble inositol hexakisphosphate. *Elife* **4**, e06074
2. Baraldi, E., Carugo, K. D., Hyvönen, M., Surdo, P. L., Riley, A. M., Potter, B. V., O'Brien, R., Ladbury, J. E., and Saraste, M. (1999) Structure of the PH domain from Bruton's tyrosine kinase in complex with inositol 1, 3, 4, 5-tetrakisphosphate. *Structure* **7**, 449-460
3. Ortega, A., Amorós, D., and De La Torre, J. G. (2011) Prediction of hydrodynamic and other solution properties of rigid proteins from atomic-and residue-level models. *Biophysical journal* **101**, 892-898
4. Schindelin, J., Arganda-Carreras, I., Frise, E., Kaynig, V., Longair, M., Pietzsch, T., Preibisch, S., Rueden, C., Saalfeld, S., Schmid, B., Tinevez, J. Y., White, D. J., Hartenstein, V., Eliceiri, K., Tomancak, P., and Cardona, A. (2012) Fiji: an open-source platform for biological-image analysis. *Nat Methods* **9**, 676-682
